# Neural encoding of socially adjusted value during competitive and hazardous foraging

**DOI:** 10.1101/2020.09.11.294058

**Authors:** Brian Silston, Toby Wise, Song Qi, Xin Sui, Peter Dayan, Dean Mobbs

## Abstract

In group foraging organisms, optimizing the conflicting demands of competitive food loss and safety is critical. We demonstrate that humans select competition avoidant and risk diluting strategies during foraging depending on socially adjusted value. We formulate a mathematically grounded quantification of socially adjusted value in foraging environments and show using multivariate fMRI analyses that socially adjusted value is encoded by mid-cingulate and ventromedial prefrontal cortices, regions that integrate value and action signals.

---

Across phylogeny, foraging decisions (e.g. patch selection, feeding behavior and duration) are strongly influenced by competitor density, food quantity and expected energy cost ^1,2^. In predation free, yet competitive, environments, avoiding competition dense patches is an adaptive strategy to maximize gain (e.g. see ^3,4^). In contrast, foraging decisions under potential threat of predation are governed by risk dilution strategies (i.e. safety in numbers) for which larger groups of conspecifics reduce the chance a particular individual will fall victim to lethal attack by predators ^5-7^. However, risk dilution strategies incur efficiency costs, reducing exploitation and harvest rates^1^. Accordingly, optimal foraging theories^1,2^ suggest that the conflicting trade-off between threat of competition and the threat of predation are mitigated by the overall fitness or socially adjusted value of the patch. Socially adjusted value, thus, is represented as the overall potential benefit, whether in the safe domain via selection of less competition dense patches or risk dilution in the threat domain via occupation of more competition dense patches and is decoupled from the observed social density of a patch. It is unclear whether humans obey these rules and whether socially adjusted value, independent of the observable statistics of a foraging environment, is represented in the human brain.

A growing body of literature is beginning to elucidate the underlying neurobiological mechanisms of foraging decision-making, albeit in the absence of threat. Human and non-human primate research has focused primarily on virtual two-patch foraging tasks, consistently highlighting regions involved in action selection and value encoding (e.g. mid-cingulate cortex [MCC] and ventromedial prefrontal cortex [vmPFC]) ^8,9^,^10^. Further, anterior to the MCC is the dorsal cingulate cortex (dACC), which has been linked to foraging decisions and the difficulty of such decisions (see for debate ^8,9^), while the vmPFC has been linked to representations of choice value during foraging tasks ^9,11^, which is in line with its role in monitoring action value, exploration and economic decisions ^12,13^. Given the functional heterogeneity of the MCC and vmPFC, one hypothesis is that these regions reflect the socially adjusted value of foraging decisions through the integration of knowledge about competition and threat, constructing a primary decision variable more immediately relevant to behavior than the simple social density of an environment. We addressed this idea by creating foraging environments that are identical except for the socially adjusted value strategies. Thus, testing conditions for both safe and hazardous environments in which the optimal strategies are inversely correlated, for example competition avoidance and risk dilution, allows us to investigate the overall socially adjusted value of the patch independent of its directly observable statistics.

In this study, participants were scanned for approximately four hours each over the course of two days while they performed a two-patch foraging task with changing level of threat and competition density. We examined multivariate, distributed neural representations involved in competition avoidance and risk dilution during a virtual foraging paradigm in which participants assessed competition density and risk of predation and made choices to enter one of two patches. First, participants learned the sequence of competitor states (specifically, the number of competitors in each patch for repeating pairs of side-by-side patches), and then were asked to select the patch in which they would like to forage. Simply selecting a patch was insufficient to receive a food reward token. Food tokens appeared at random times and locations in the patch, and the player was required to navigate to it before other competitors in the patch. Therefore, the higher the density of competitors, the less likely it was that the subject would be able to acquire the token. However, in some patches (identified by the color), a predator could appear randomly at any time or location and would capture the player or one of the other competitors, also at random.

We hypothesized that subjects would adapt their foraging decisions based on the underlying socially adjusted value of a patch, being higher in low social density patches during safe foraging (as a result of high reward availability), and higher in more socially dense patches during threat of predation (as a result of decreased risk of predation). We additionally hypothesized that these decisions would be supported by neural encoding of socially adjusted value independently of social density per se. Importantly, our safe and hazardous patches were matched for the effort of decision, energy costs, and competition and reward density. Further, patch switch costs were zero, therefore allowing us to investigate pure contextual changes in socially adjusted value of the decision.

We assessed decision making by condition by computing the percentage of trials in which the player selected the patch with fewer competitors (Fig. 1, panel E.), and in more fine grained detail by calculating a difference score reflecting the number of competitors present in each patch at the time of decision. Participants selected the less populated patch in 89% of decisions for the safe condition and in 32% of decisions in the threat condition, inclusive of all trial durations (χ^2^ = 4046, p<.0005, proportion; t(21) = 9.07; p<.0005, total count). Further, we observed different average difference scores for safe and threat conditions (t(24) = 9.14, p<.0005), indicating a context dependent shift in behavioral decision-making based on socially adjusted value of each patch.

**Figure 1.**
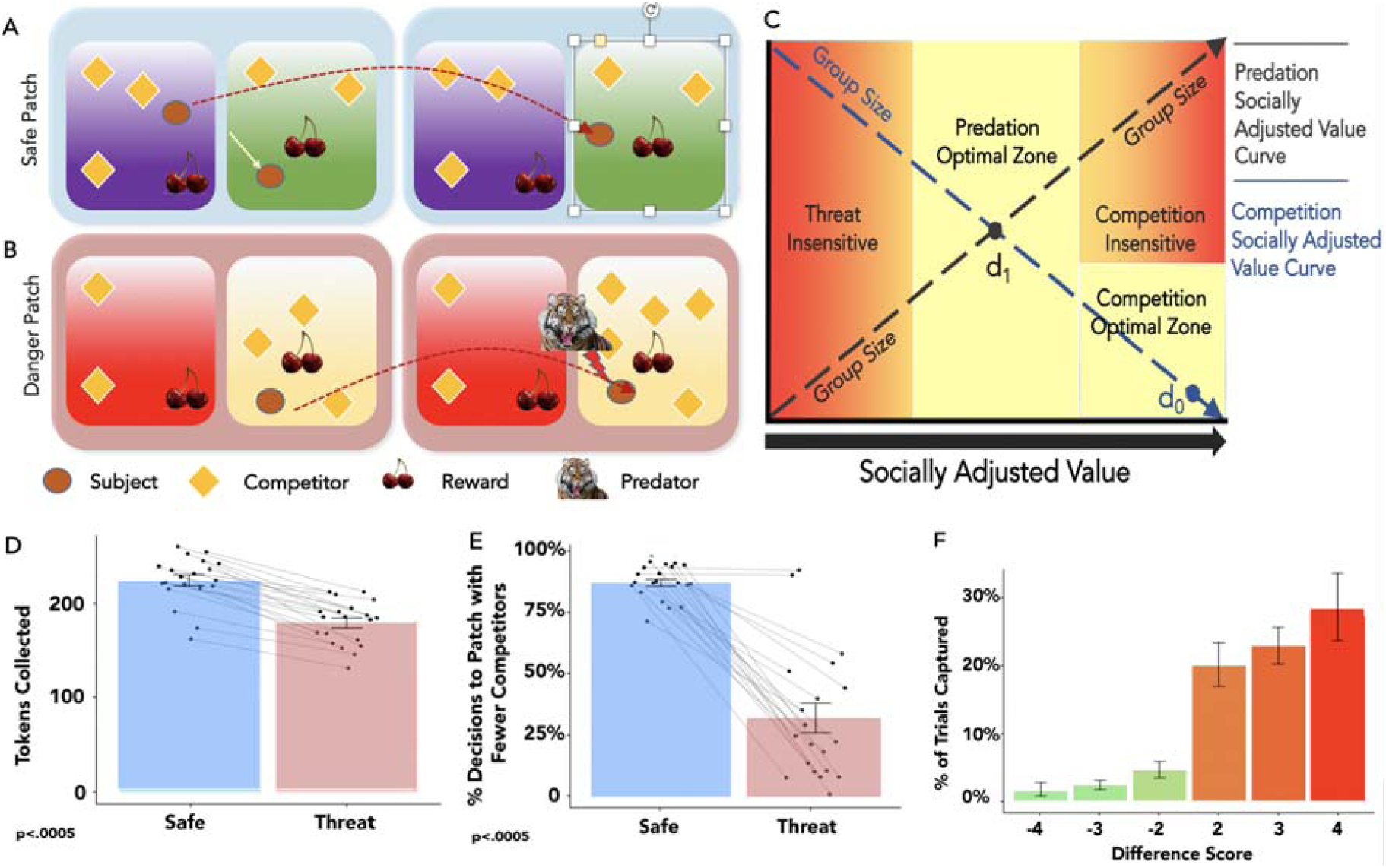
A: Task Design. In the safe play phase the player selects the desired patch and then competes to capture rewards; B: in the danger phase the player selects the desired patch and then competes for rewards, and is subject to potential capture by the predator. In the safe condition the trial ends after time expires. The side-by-side patches were the same color during the experiment and are just color-coded here to clarify the differences between safe and threat conditions. C: The risk calculus that informs individual decision making based on socially adjusted value. In safe patches (blue line) the optimal strategy is to select the patch with the fewest number of competitors, thus maximizing reward gain and socially adjusted value. In dangerous patches (black line) increasing group size threatens the ability to capture rewards but dilutes risk of being the target of the predator. d_o_ represents a decision in which socially adjusted value has been maximized in the safe condition; d_1_ represents an individual with moderate risk tolerance in the danger condition, willing to select a patch with several competitors in order to reduce capture risk while still competing for rewards; D: Players collected significantly more rewards in safe than in threat patch configurations; E: Players spontaneously adopted a competition avoidance strategy in the safe condition, while most but not all players adopted a risk dilution strategy in the threat condition; F: Probability distribution of being captured as a function of the difference score on a given trial. A large positive difference score indicates presence in a patch with few or one other competitor. A large negative score indicates presence in a patch with several other competitors.

The foregoing tendency to congregate in numbers in the presence of potential predation optimized the probability of avoiding capture, even for the smallest patch difference (Figure 1, panel F). The converse was true for the safe context, such that avoiding competition resulted in the maximizing token collection (Figure 1, panel D). Decisions to forage in patches with fewer competitors increased capture probability compared with foraging in patches with more competitors, even for the closest value differences, e.g. between having 2 more players in the patch or two fewer players in the patch (χ^2^=109.9, p<.0005). Players also collected significantly fewer food tokens in the threat than the safe condition, likely reflecting both increased competition and distraction due to anticipation of the predator (t(38) = 5.93, p<.0005; Fig 1, panel D). To assess the effect of distraction of threat above and beyond basic competition we assessed mean token collection at each level of competition across both safe and threat conditions, and overall across conditions (see Fig S2 in supplementary materials).]

Optimization in the safe domain via selection of less competitive patches held across all four blocks and began early in the first block, reflecting spontaneous and swift acquisition of adaptive decision-making behavior. Likewise, most, but not all participants quickly adopted a risk dilution strategy in the presence of predation, evidenced by consistent decision behavior across all four trial blocks. While some participants adopted an identical strategy across trial type (e.g. safe / threat), most decisions in safe trials reflected a competition avoidance strategy, that is, to a patch with fewer competitors in order to maximize gain (see Figure 1, panel A), while the opposite was true for threat trials, on average, suggesting a risk dilution strategy (see Figure 1, panel B).

In order to provide a mathematically grounded quantification of socially adjusted value, the key decision variable emerging in our behavioral analyses, we fit a model to subjects’ decisions that made decisions based on the socially adjusted value of each patch (see Methods). Socially adjusted value depended on the average number of points collected in each condition, and included a free parameter representing the value of receiving a shock, which was negative for all but one subject (Figure 2A). As expected, inferred socially adjusted value from the model decreased with more competitors during safety and increased during threat (Figure 2B), and the difference in socially adjusted value between the two patches was strongly predictive of choice (Figure 2C), correctly predicting 66.72% of choices. Predicted probabilities based on socially adjusted value difference were also well calibrated with respect to true choice probabilities (Figure 2D).

**Figure 2.**
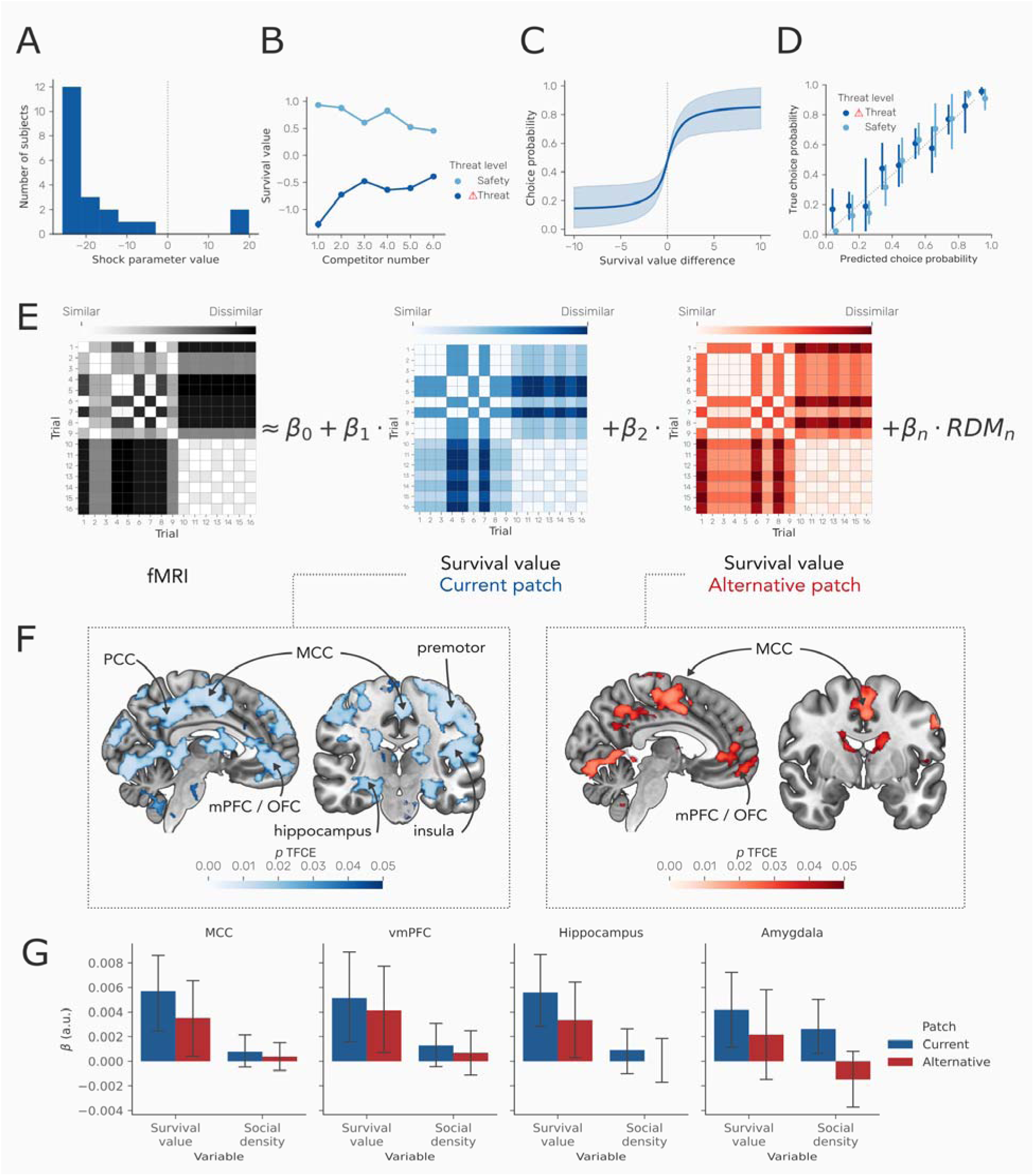
A: Values of the shock cost parameter from our behavioral model, indicating that shocks were perceived as a cost for all but one subject. B: Socially adjusted value across task conditions, demonstrating that socially adjusted value depends on both threat level and the number of competitors. C) The probability of choosing a patch based on the difference between its socially adjusted value and that of the alternative patch. The function shown represents the results of logistic regression models predicting choice from socially adjusted value difference (see methods). D) Calibration plot showing predicted probability of choice based on socially adjusted value difference derived from a logistic regression model versus the true choice probability for safe and threat conditions. Probabilities are binned into 10% bins, and error bars represent 95% confidence intervals across subjects. E: Representational dissimilarity matrices (RDMs) for the neural data (hypothetical example shown left) and task conditions of interest (right). Neural RDMs were modeled as a function of task RDMs using linear regression, with each RDM weighted by a weight parameter β_RDM_. Here, RDMs are shown for a subset of trials for a single subject. In the task RDMs, rows and columns represent individual trials, while the colors in the matrix represent the difference in socially adjusted value between that trial and other trials. RDMs are shown for the socially adjusted value of the current and alternative patch, and other RDMs are described the methods and results. F: Left panel: effects of the current patch socially adjusted value RDM, showing widespread effects, including the mid-cingulate cortex (MCC), posterior cingulate cortex (PCC), medial prefrontal cortex (mPFC), orbitofrontal cortex (OFC).. Right panel: effects of the alternative patch socially adjusted value RDM, showing effects across similar areas, with the most prominent clusters in the mid cingulate cortex (MCC), and mPFC/OFC. Maps represent p values determined using threshold-free cluster correction (TFCE), thresholded at p < .05. G) Extracted beta values from the MCC and vmPFC, taken from the AAL atlas ^14^ mid-cingulate and frontal medial orbital regions respectively, in addition to the AAL hippocampus and amygdala regions. Higher values indicate greater similarity between the task RDM and the neural RDM. These values are provided for illustration only; significance was determined using voxelwise tests as shown in (F). Error bars represent 95% confidence intervals.

We next sought to determine how decision variables are encoded in the brain. We used representational similarity analysis (RSA) to identify regions where activity patterns aligned with the task structure during the decision-making phase of the task, during which subjects were aware of available options but could not yet make a motor response. Critically, RSA allowed us to identify multivariate representations of key decision variables, rather than assessing changes in single-voxel activity levels as in traditional univariate analyses, through identifying regions where neural similarity across conditions aligns with similarity in the properties of the task being performed. While univariate analyses could only determine whether socially adjusted value is associated with activity in each region, RSA allows us to determine whether the multivariate representation of socially adjusted value in a region is consistent across different conditions. We computed representational dissimilarity matrices (RDMs) for BOLD responses to each trial and used linear regression to predict the observed pattern of neural similarity from RDMs for task conditions (Figure 2D). First, we computed task RDMs based socially adjusted value (as determined using our behavioral model) in the current patch (the patch selected on the previous trial) and the alternative. Second, we computed RDMs representing the socially adjusted value of the current and alternative patch. We also included RDMs representing the difference in competitors and socially adjusted value between patches and included an RDM representing the effect of threat.

We used a searchlight approach, performing the RSA analysis in within 6mm spheres across the brain to localize regions encoding task-relevant representations. This identified a distributed network of regions encoding the socially adjusted value of the alternative patch during decision-making, including the MCC, posterior cingulate cortex (PCC), medial prefrontal cortex (mPFC), and orbitofrontal cortex (OFC) (voxel ps < .05, threshold-free cluster correction, Figure 2E). The socially adjusted value of the current patch was also represented in these regions, but was additionally represented in the premotor cortex, hippocampus, and anterior insular cortex (voxel ps < .05, threshold-free cluster correction, Figure 2E). Importantly, our analysis approach isolated unique effects of each task RDM, controlling for effects of other task RDMs. In contrast, the number of competitors in the alternative patch was only represented in the left pre-supplementary motor area (voxel ps < .05, threshold-free cluster correction), and no area represented the number of competitors in the current patch. Notably, we found no region encoding the difference between patches, either in terms of number of competitors or socially adjusted value. Threat level (i.e. safe or at risk of predation) was encoded in a wide range of cortical and subcortical regions (see Figure S3), including the MCC, vmPFC, hippocampus and amygdala.

Our results demonstrate that humans adaptively select social environments based on their socially adjusted value, using a competition-avoidant strategy when threat is absent and switching to a risk-diluting strategy in the presence of threat. Importantly, the key variable underpinning this decision, the socially adjusted value of a foraging patch, was encoded in a network focused on the MCC and vmPFC, suggesting that these regions integrate information about social density and threat to calculate the overall socially adjusted value of both the current patch and an alternative. Our results identify distributed neural systems representing key decision variables underlying adaptative foraging in response to competition and threat.

Behaviorally, we found that subjects adapted their decision-making strategy based on the socially adjusted value of foraging patches. Under conditions of safety subjects were biased towards patches with low competition density, as found in previous work ^10^, representing a competition-avoidant strategy. In contrast, when under threat, subjects chose patches with high social density, representing a risk-dilution strategy. These results indicate that while foraging in social environments with threat of predation, human subjects base their decisions on the socially adjusted value of available patches. In the absence of predation risk, immediate survival depends on maximizing food resources, and is therefore highest in environments with the fewest competitors. When under threat of predation, survival focus shifts to risk of capture, and is higher in environments with a greater number of competitors, and remains so until competition density outweighs the risk dilution benefit. In real-world environments, socially adjusted value will depend on the interaction between multiple factors, and our task presents a simplistic case where reward is readily available. For example, when food is persistently scarce, socially adjusted value is likely to be relatively high in the absence of competition even under threat.

At the neural level, we found that the socially adjusted value of a social decision environment was encoded across a distributed network of regions, primarily located in the vmPFC, OFC, MCC and PCC. In contrast, we found little evidence for a representation of the number of competitors per se, or the difference in socially adjusted value or social density between patches. These results indicate that when making decisions about foraging patches in the presence of competitors, the brain predominantly represents the socially adjusted value of a potential patches, regardless of whether this represents risk of predation or risk of competition for food, while potentially disregarding the observed number of competitors, or the difference between available patches. Importantly, this suggests that in social foraging environments the brain is representing the value of social density in terms of a socially adjusted value, rather than simply representing the numbers of competitors, and suggests that the brain encodes the value of independent options rather than their difference. Our use of RSA allowed us to focus on the multivariate representations of these variables in the brain, rather than relying on single-voxel activity changes. Thus, the regions we identify do not simply show similar activity levels across high socially adjusted value conditions but represent survival in the same manner across conditions. Our focus on multivariate representations of decision variables during foraging, as opposed to overall activity levels, distinguishes this work from prior studies, which to date have exclusively used univariate approaches. Thus, while activity levels may be modulated by the difference between options, the pattern of activity represents the value of each option independently.

Concerning socially adjusted value, two regions are of historical interest ^9^, the MCC and vmPFC. These regions are active across both current and alternative patches. Importantly, our results show that these regions encode the socially adjusted value of an environment across safe and threat conditions, rather than the social density of competition alone. What are these two regions computing? It is important to state that our MCC cluster is more posterior than the dACC area that has been linked to the difficulty of foraging decisions and the value of alternative options^8,10,13^. Our task conditions of interest are largely orthogonal to decision difficulty. Thus, our lack of dACC activity may reflect the matching of difficulty across conditions. Our results also indicate that the MCC is not purely signaling the value of an alternative option, or the difference between options, but simultaneously represents both the value of both the current and alternative option.^9,11,15^ However, based on the knowledge of this region’s connectivity and function, which suggests it plays little role in value-based, goal-directed behavior *per se*^16^, it seems incorrect to say that the MCC reflects pure value. Instead, the MCC may act as a hub, coordinating emotional responses and motor actions according learned values^17^, particularly when under threat^18^. Of note, threat level was also encoded in these regions, suggesting that they represent predation threat in addition to decision variables dependent on threat.

At face value, our results are in line with the role of the vmPFC in valuation of options during foraging. In prior work using foraging paradigms, however, the vmPFC has been implicated as a comparator of options, while our work suggests that it independently signals the values of multiple options ^19^. Others have shown that damage to the vmPFC can result in poorer learning of the value of spatial locations of rewards, supporting the role of the vmPFC in domain-general valuation^11,20^. While more novel task designs are required to dissociate their roles precisely, the extensive prior work on the roles of the vmPFC and MCC suggests that they are perfectly situated to calculate different aspects of foraging decisions based on socially adjusted value. While the vmPFC appears important in representing the socially adjusted value of options itself, potentially facilitating the decision-making process, MCC is likely involved in the coordination of motor actions (i.e. to stay or go). This is particularly important in the current study as the socially adjusted value encompasses both competition avoidance and risk dilution strategies.

## Supplementary Materials

### Methods

#### Subjects

Participants were screened for requirements to participate, including standard health measures that determine inclusion for fMRI experiments. Following the screening, 22 (6F; mean age: 31; range: 18-49) participants were trained on and completed a computerized, virtual foraging task featuring two conditions: safe and threat (Figure 1). Behavioral and neural data were lost for one session for one participant and neural data for another participant resulting in reporting of 20 participants for behavioral and neural data analyses. Participants were scanned for approximately four hours each over the course of two days. The threat condition involved the possibility of electric shock, administered to the underside of the left wrist. Electric shock intensity was individually calibrated prior to the task to an aversive, yet tolerable level.

#### Task

The task was a dynamic foraging game designed to investigate the effects of competition and threat on foraging decisions and behavior. Participants foraged for two, 45 minute blocks per day over two days for a total of four blocks (Figure S1) and approximately four hours of total scanning including structural scans.. Each block maintained an identical trial structure, the difference being the number and cycle of competitors in the two patches. During each block a consistent cycle of competitors repeated, such that the player could quickly learn to predict the configuration (e.g. number of competitors in each patch) of the following part of the cycle. The game consisted of two patches containing 1-6 other AI players that foraged the environment for rewards, which appeared on a consistent, periodic basis.

#### Cycles

Each session was characterized by a regular progression of cycles (Figure S1). Each cycle consisted of three distinct competitor states that repeated sequentially throughout a session. During session one, the left patch (denoted P1) progressed in such a manner that if the participant saw one competitor during the current trial, on the following trial the participant would see five competitors, followed by two competitors. The cycle repeated for the duration of the session. The same mechanism operated for the right patch (denoted P2) such that when the participant saw one competitor in the left patch (P1), four would be present in right patch (P2); if five competitors were present in the left patch (P1), then one competitor would be present in the right patch (P2); if two competitors were present in the left patch (P1), then five competitors would be present in the right patch (P2). Different, repeating competitor state progressions were used for sessions two, three and four (Figure S1).

#### Trial Types

Trial types included a short duration, immediate decision (SI) in which participants were briefly shown the upcoming competitor state, prompted to select either the left (P1) or right (P2) patch, were placed in the selected patch and foraged for seven seconds; short duration, later decision (SL) in which participants were briefly shown the current competitor state, prompted to select either the left (P1) or right (P2) patch for the current state in the cycle, or, if they wished, the next state in the cycle, were placed in the selected patch and foraged for seven seconds; and long duration, immediate decision (LI) in which participants were briefly shown the current competitor state, prompted to select either the left (P1) or right (P2) patch for the current state in the cycle, were placed in the selected patch, and foraged for 14 seconds. During the 14 second period the patch state would change once (e.g. every 7 seconds). Importantly, LI trials were single condition trials such that if the first 7 seconds was safe, the latter portion of the trial was also the safe condition, and if the first 7 seconds was under threat, the latter portion of the trial was also the threat condition. Since few subjects utilized the later decision option, data analyzed includes only immediate decisions categorized as SI and LI. In each of these instances the trial format was identical, and participants selected from among the patch options displayed to them prior to engaging in foraging activity, enabling us to collapse decisions across trial types.

#### Decision

At the beginning of each trial, two patches were displayed for three seconds, each with a different number of competitors (a competitor state). After the three second period elapsed the patches disappeared, and the participant had three seconds to make a decision regarding the patch in which they would like to forage. In the SI condition, the participant chose between the two patches visually displayed, and was placed in the selected patch for the remainder of the trial. In the SL condition, the participant could choose either from among the patches visually displayed or the next iteration of patches per the repeating competitor states. If the player selected a patch from the visual display the trial would proceed identical to the SI trial type; however, if instead a patch from the next state in the cycle (not visually displayed) was selected the patches would immediately change to the following competitor configuration state and the player would proceed to forage for the remainder of the trial. In the LI condition, the patch configuration iterates halfway through the trial, forcing the player to consider both the current and next state in the cycle as they make a decision from among the visually displayed patches.

#### Gameplay

After choosing a patch, participants foraged using a diamond button box with four easy to use buttons indicating directions up, down, left and right, until trial ended, which occurred due to a time parameter or appearance of a virtual predator. AI competitors were opaque (the player could not move through AI competitors and vice versa) and programmed to chase food tokens within a predetermined radius at the same speed as the player. During or after the threat trials the game paused for two seconds when the virtual predator appeared, after which the predator moved towards and captured a player at random, ending the trial. The predator was more likely to attack the patch in which the participant was located, but not more likely to attack the participant than other players. If the participant was captured they received an electric shock and lost a small portion of the accumulated earnings acquired over the course of the block.

Safe trials consisted of making decisions based on the configuration of competition in each patch and the learned repeating cycles that enable recognition of the location within a cycle and hence, the upcoming configuration. Threat trials included the appearance of a virtual predator, adding a relevant parameter to participants’ decisions. Selecting a patch with few players increases risk of capture by a predator, but also increases the chance of acquiring rewards. Occupying a patch with several other players dilutes risk of capture by a predator, but decreases the chance of acquiring rewards.

**Figure S1.**
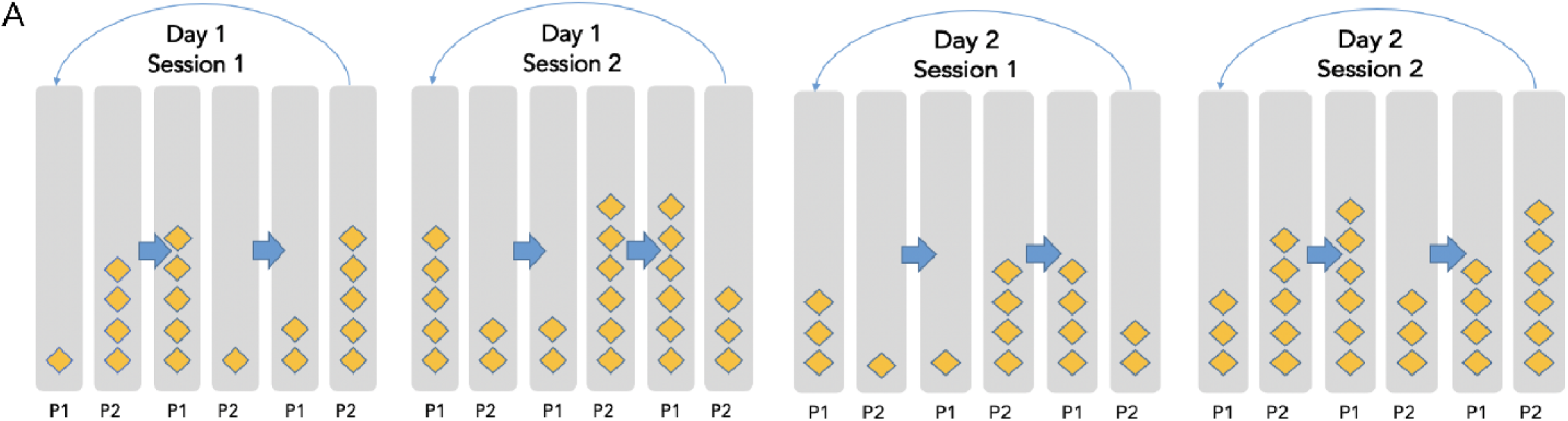
Task Design Competitor States and Cycles. The design featured two sessions each day over two days, in which patch configurations changed in repeating cycles; each diamond represents a competitor such that, for example, on Day 1 Session the first patch state included one competitor in P1 and four competitors in patch 2; P1 (patch 1); P2 (patch 2), followed by a state in which P1 contained five competitors and P2 contained one competitor, followed by a state in which P1 contained two competitors and P2 contained five competitors.

Each block was divided into eight (8) sub blocks consisting of 18 trials of which half were safe trials and half were threat trials. Based on this structure, participants completed a total of 576 trials, balanced between the three trial types detailed above (SI, SL, LI). Threat trials ended when a virtual predator appeared and captured a player. The task duration over the course of two days was approximately 3.5 hours, including a short training period. During the training period the participant observed changing patch configurations as a fixed 3-change cycle was repeated. For example, the left patch might cycle through the following number of competitors: 6, 4, 3 such that if the player observes four players in the patch, the next cycle will display three players with 100% certainty. Participants completed a post task questionnaire, were debriefed on the experimental purpose, and paid $120 for participation.

#### Behavioral data acquisition and analysis

Participants’ x-y coordinates were recorded at a sampling rate of 30Hz. Other variables collected included reward spawn location and collection; patch selection decisions; safe / threat value of each environment, number of competitors in each environment, captures and shocks received.

Decisions tallies were generated for two focal variables including whether 1) the player selected the patch with fewer competitors; 2) the player selected a patch currently observed or a prospective patch. These decisions were analyzed based on several factors including 1) the trial type, e.g. safe or threat; 2) the trial length, e.g. short or long; 3) the block, e.g. low competitor numbers or high competitor numbers; and 4) the actual number of competitors in each patch.

#### Behavioral modelling

To quantify socially adjusted value at the individual subject level, we fit a model to the behavioral data. This model represented decisions as being based on the socially adjusted value of the two potential patches. Socially adjusted value was determined based on the average accumulated points across the task in each condition (i.e. every combination of competitor number and threat level) across the entire task, accounting for losses incurred when caught, and was calculated independently for each subject. In addition, we included the probability of being caught in each condition, multiplied by a free parameter representing the cost of receiving a shock. Thus, socially adjusted value was dependent on both the number of points collected and the cost of shocks. Notably, the mean number of tokens collected across competitor number and condition favored the safe condition, except at a value of three competitors at which no difference in mean token collection was evident (Fig S2), suggesting an effect of threat above and beyond competition vis a vis socially adjusted value. The socially adjusted value of condition X was therefore calculated as:

A softmax function with a free temperature parameter (was used to transform the values of the two patch options (A and B) to choice probabilities. This calculation used only the value of the two patches shown on screen, rather than later patches on LI trials, as behavioral results demonstrated little evidence for subjects considering these later patches.

Models were fit in Python using PyMC3, using MCMC sampling with 4 chains of 2000 samples. For further analyses, we used the mean of the posterior distribution over parameters.

**Figure S2.**
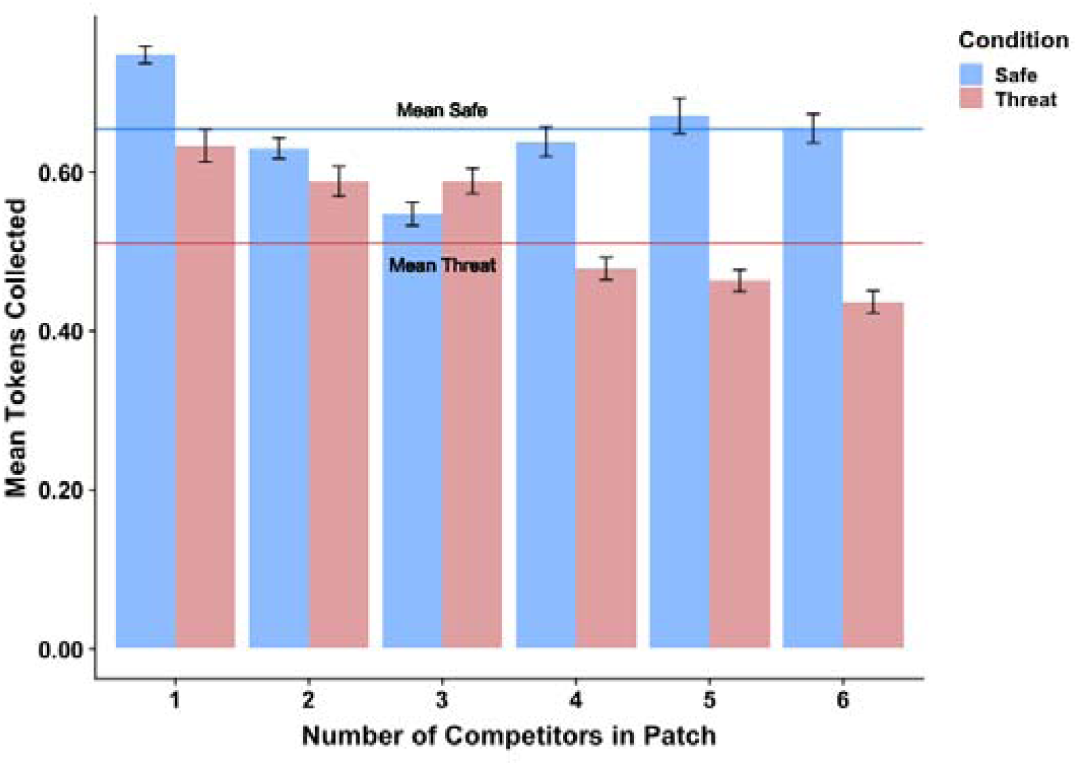
Mean token collection across competitor number in selected patch across conditions. The average difference favored the safe condition, especially in patches with higher numbers of competitors. Dashed line represents the absolute difference of mean values. Only patches with three competitors showed no difference in mean token collection across conditions.

#### FMRI data acquisition

fMRI data was collected using a 3T Prisma scanner in the Caltech Brain Imaging Center (Pasadena, CA) with a 32-channel head receive array. BOLD contrast images will be acquired using a single-shot, multiband T2*-weighted echo planar imaging sequence with the following parameters: TR/TE = 1000/30 ms, Flip Angle = 60°, 72 slices, slice angulation = 20° to transverse, multiband acceleration = 6, no in-plane acceleration, 3/4 partial Fourier acquisition, slice thickness/gap = 2.0/0.0 mm, FOV = 192 mm × 192 mm, matrix = 96 × 96). Anatomical reference imaging will employ 0.9 mm isotropic resolution 3D T1w MEMP-RAGE (TR/TI/TE = 2550/1150/1.3, 3.1, 4.0, 6.9 ms, FOV = 230 m x 230 mm) and 3D T2w SPACE sequences (TR/TE = 3200/564 ms, FOV = 230 mm x 230 mm). Participants viewed the screen via a mirror mounted on the head coil, and a pillow and foam cushions were placed inside the coil to minimize head movement. Electric stimulation was delivered using a BIOPAC STM100C.

#### FMRI preprocessing and data analysis

Participants’ data were preprocessed using fMRIprep ^21^, an automated preprocessing pipeline consisting of bias correction, registration to a standard template, slice timing correction, and motion correction. Noise timeseries, derived from white matter and cerebrospinal fluid (CSF) masks, were extracted using aCompCor ^22^.

First level models were constructed using FSL 6.0 ^23^. Each run of the task was modelled and estimated separately, including single regressors for each task periods of no interest. Data was normalized such that each voxel had a mean value of 100 prior to analysis. As our analyses focused on the decision period of the task, we modelled this period of each trial using a separate regressor, resulting in independent beta maps for each trial ^24^. We also included framewise displacement, six motion parameters, white matter and CSF timeseries as regressors of no interest to account for signals related to motion and physiological processes. As the number of trials per condition was dependent on subjects’ choices, we excluded subjects who did not experience the +/-2 or 3 competitor difference conditions. This resulted in the exclusion of a single subject.

Maps for each condition were subsequently used for representational similarity analysis (RSA) using a searchlight approach. This involved calculating representation dissimilarity matrices (RDMs) for the neural data by computing the Spearman correlation distance between voxel-level representations within a 6mm sphere for each condition across the entire task. This resulted in RDMs with as many rows and columns as trials in the task. To quantify associations between the neural RDMs and task variables, we calculated task RDMs representing the distance between conditions in terms of task features of interest.

We focused on two RDMs representing task features of particular interest: First, the number of competitors in each patch, with one RDM for the current patch, one for the alternative, and one for the difference between the two. Second, the socially adjusted value of each patch, also including RDMs for current, alternative, and difference between patches. We then used linear regression to determine the influence of task RDMs on neural RDMs, with each RDM being weighted by its own β parameter. We also included a further RDM representing threat level, along with RDMs representing condition similarity in run number (where conditions from the same run have highest similarity) and session number (where conditions from the same session have highest similarity). This produced whole-brain maps representing the β weights for the RDMs of interest, with each voxel representing the effect in the 6mm sphere of which it was the center.

Maps from the searchlight RSA analysis were thresholded using the randomize function in FSL, using a one-sample t-test against zero. Statistical significance at every voxel was determined using threshold-free cluster correction with 5000 permutations and 10mm variance smoothing.

**Figure S3.**
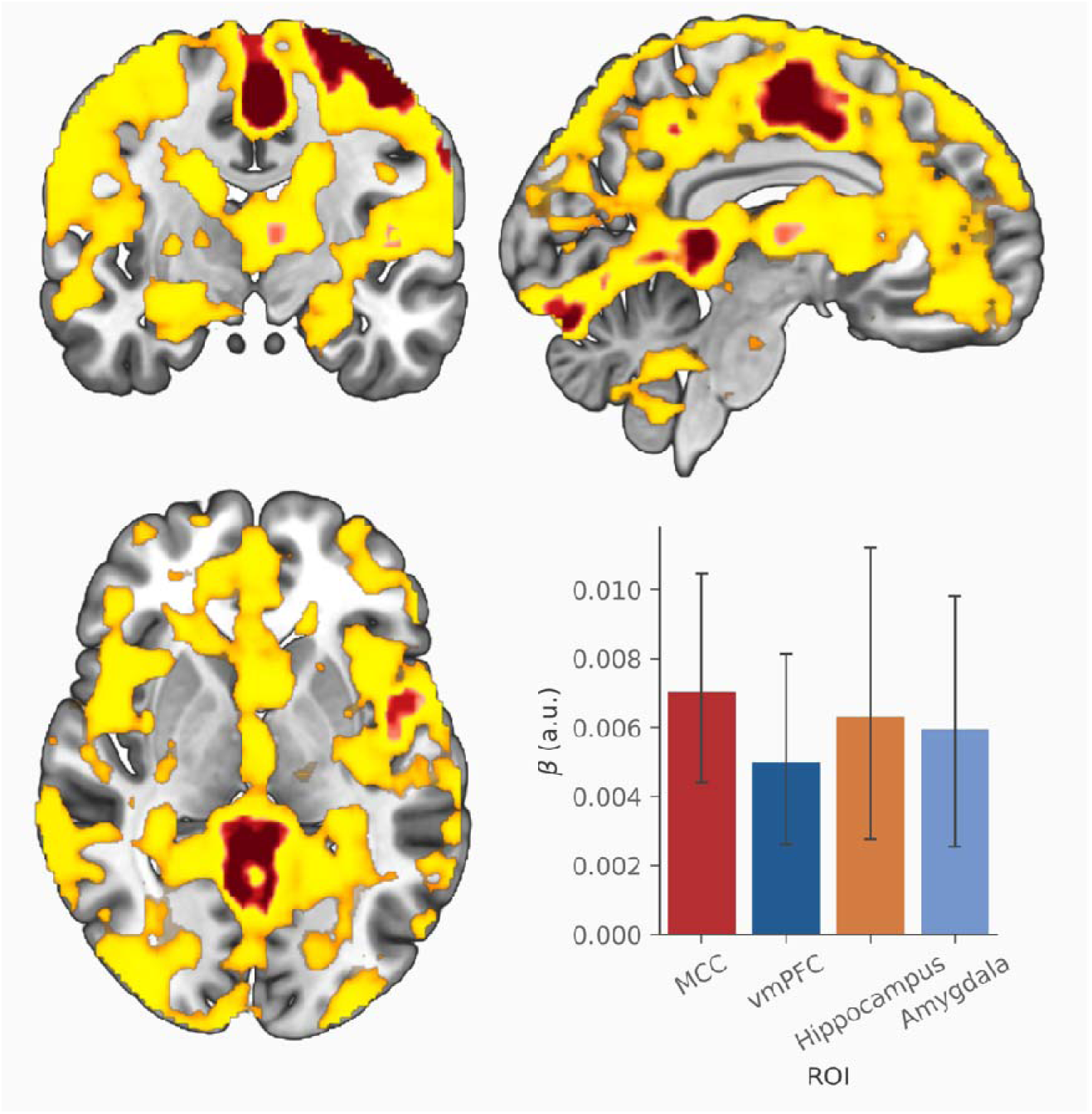
Regions representing threat versus safety in the RSA analysis. The map represents p values determined using threshold-free cluster correction (TFCE). Yellow/orange shows areas where p < .05, while red shows areas where p < .001. The bar plot represents values from the mid-cingulate cortex (MCC), ventromedial prefrontal cortex (vmPFC), hippocampus and amygdala. Error bars represent 95% confidence intervals across subjects.

## Notes

### Competing Interest Statement

The authors have declared no competing interest.

